# SharpTNI: Counting and Sampling Parsimonious Transmission Networks under a Weak Bottleneck

**DOI:** 10.1101/842237

**Authors:** Palash Sashittal, Mohammed El-Kebir

**Affiliations:** Department of Aerospace Engineering, University of Illinois at Urbana-Champaign, Champaign, IL 61801, USA; Department of Computer Science, University of Illinois at Urbana-Champaign, Champaign, IL 61801, USA

**Keywords:** Phylogenetics, Phylodynamics, Phylogeography, Migration, Transmission, Infection, Outbreak, Approximate counting, Almost-uniform sampling, Satisfiability

## Abstract

**Background:** Technological advances in genomic sequencing are facilitating the reconstruction of transmission histories during outbreaks in the fight against infectious diseases. However, accurate disease transmission inference using this data is hindered by a number of challenges due to within-host pathogen diversity and weak transmission bottlenecks, where multiple genetically-distinct pathogenic strains co-transmit.

**Results:** We formulate a combinatorial optimization problem for transmission network inference under a weak bottleneck from a given timed phylogeny and establish hardness results. We present SharpTNI, a method to approximately count and almost uniformly sample from the solution space. Using simulated data, we show that SharpTNI accurately quantifies and uniformly samples from the solution space of parsimonious transmission networks, scaling to large datasets. We demonstrate that SharpTNI identifies co-transmissions during the 2014 Ebola outbreak that are corroborated by epidemiological information collected by previous studies.

**Conclusions:** Accounting for weak transmission bottlenecks is crucial for accurate inference of transmission histories during outbreaks. SharpTNI is a parsimony-based method to reconstruct transmission networks for diseases with long incubation times and large inocula given timed phylogenies. The model and theoretical work of this paper pave the way for novel maximum likelihood methods to co-estimate timed phylogenies and transmission networks under a weak bottleneck.

## Background

Accurate inference of the transmission history of an infectious disease outbreak is pivotal for real-time outbreak management, public health policies and devising disease control strategies for future outbreaks [1]. Traditional epidemiological approaches are fieldwork intensive and aim to uncover contact histories and exposure times of hosts to disease sources. With decreasing costs of genomic sequencing, molecular epidemiology has complemented these traditional approaches to effectively analyze and manage disease outbreaks.

Given genomic and epidemiological data, the key challenge is to infer the evolutionary history of the pathogen isolates and the transmission history of the hosts. Importantly, while the phylogeny of the pathogen isolates captures the evolutionary history of the outbreak, it does not necessarily match the transmission history of the outbreak [2]—this mutation-migration discordance also arises in metastatic cancers [3]. In particular, methods that assume that transmission events coincide with branching events in the phylogeny are only applicable in the context of pathogens with low mutation rates, short incubation times and acute infections [4–8]. By contrast, pathogens with high mutation rates and long incubation times lead to *within-host diversity*. This diversity is either the result of infection by multiple strains or arose after infection by a single strain. Most current methods assume the latter, an assumption known as a *complete transmission bottleneck* [9–14].

Under a *weak transmission bottleneck* multiple genetically-distinct strains of the pathogen are simultaneously transmitted from a donor to a recipient through a non-negligibly small inoculum. Large inoculum sizes have been observed in a number of diseases [15]. There are two recent methods that partially support a weak transmission bottleneck [16,17]. While SCOTTI allows a single host to be infected by multiple strains, it does not support the simultaneous transmission of these strains and considers each in isolation [16]. On the other hand, BadTrIP supports simultaneous transmission but does so only at single locus resolution rather than genome scale [17]. Supplementary Table S1 provides a summary of current methods.

Here, we formulate the Transmission Network Inference (TNI) problem under a weak bottleneck for a given timed phylogeny (Fig. 1). In this problem, we use the principle of parsimony to minimize the number of co-transmissions, which each may comprise of multiple transmitted strains. We prove hardness for the optimization and sampling versions of the problem. We introduce SharpTNI, a method to uniformly sample optimal solutions and quantify the size of the solution space. On simulated data, we show that SharpTNI accurately counts and samples parsimonious transmission networks, scaling to large datasets. We analyze a dataset from the 2014 Ebola outbreak [18], showing that SharpTNI outperforms SCOTTI and recapitulates previously documented co-transmissions.

**Figure 1.**
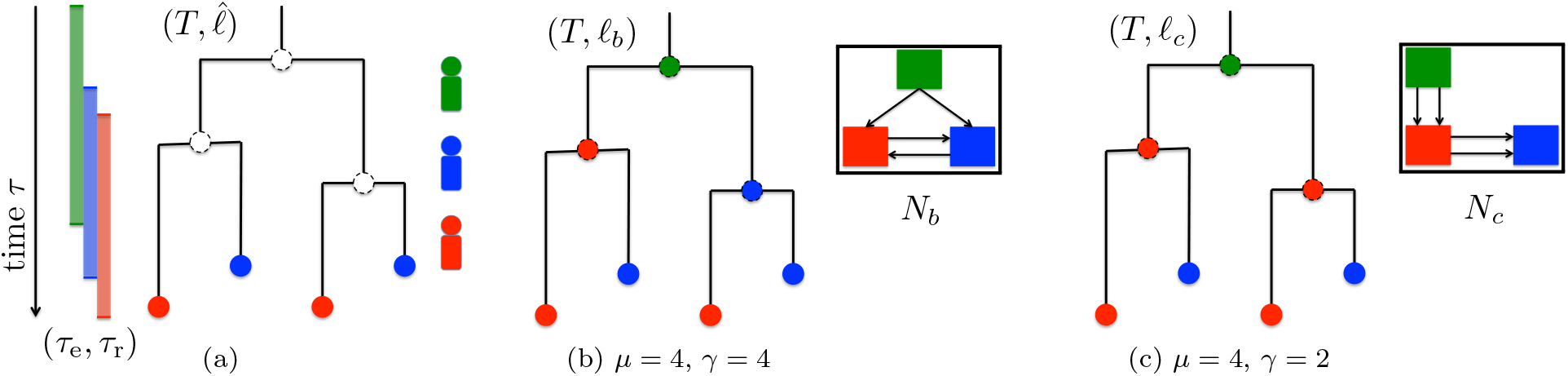
Overview of the Transmission Network Inference (TNI) problem. (a) The evolutionary history of the pathogenic strains in an outbreak is described by a timed phylogeny *T*, assigning a time-stamp *τ*(*v*) to every vertex *v*. In addition, each leaf *v* is labeled by the host 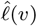 where the corresponding strain was observed (indicated by colors). Epidemiological data further constrain the entrance and removal time [*τ*_e_(*s*); *τ*_r_(*s*)] of each host *s*. In the TNI problem, we seek a host labeling *ℓ* with minimum transmission number *μ* and subsequently smallest co-transmission number *γ*. (b) Host labeling *ℓ*_*b*_ with minimum transmission *μ*\* = 4 but not the smallest co-transmission number *γ* = 4, resulting in a complex transmission network *N*_*b*_. (c) Host labeling *γ*_c_ with minimum transmission *μ*\* = 4 and smallest co-transmission number *γ*\* = 2, resulting in a parsimonious transmission network *N*_*c*_. Supplementary Fig. S1 shows all 9 minimum-transmission host labelings.

## Results

This section outlines the problem statement, the complexity results and the results obtained by applying our method SharpTNI to simulated and real datasets.

### Problem Statement

Let *T* be a tree rooted at vertex *r*(*T*) with vertex set *V*(*T*), leaf set *L*(*T*) and edge set *E*(*T*). We denote the children of a vertex *u* by *δ*_*T*_ (*u*). Conversely, the unique parent of a non-root vertex *u* ≠ *r*(*T*) is denoted by *π*_*T*_ (*u*). We write 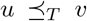 if vertex *u* is ancestral to vertex *v*, *i.e*. vertex *u* is present on the unique path from *r*(*T*) to vertex *v*. Note that 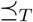 is reflexive. We say that *u* and *v* are incomparable if neither 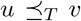 nor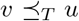 holds. We omit the subscript *T* from 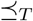, *δ*_*T*_ and *π*_*T*_ if it is clear from context. We denote the subtree of *T* rooted at vertex *v* by *T*_*v*_.

Give a set Σ of hosts, the key objects in this paper are *a timed phylogeny T* and *host labeling ℓ* : *V* (*T*) → Σ, which are defined as follows.

#### Definition 1

*A* timed phylogeny *is a rooted tree T whose vertices are labeled by time-stamps τ* : *V* (*T*) → ℝ^≥0^ *such that τ* (*u*) < *τ* (*v*) *for all pairs u*, *v of vertices where* 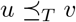.

#### Definition 2

*A* host labeling *of a timed phylogeny T is a function ℓ* : *V* (*T*) → Σ, *assigning a host ℓ*(*u*) *to each vertex u of T*.

Intuitively, time moves forward when traversing down a timed phylogeny *T* starting from the root *r*(*T*). A leaf *u* of *T* corresponds to a strain that has been removed from the population at time *τ* (*u*), due to treatment or death of the corresponding host *ℓ*(*u*). On the other hand, an internal vertex *u* of *T* corresponds to a strain that infected host *ℓ*(*u*) at time *τ* (*u*).

A timed phylogeny *T* combined with a host labeling *ℓ* constrains the set of allowed transmissions in the following three ways. First, an edge (*u v*) of *T* is a *transmission edge* if *ℓ*(*u*) ≠ *ℓ*(*v*). Second, a *transmission event* Ψ is a subset of transmission edges between the same pair of hosts that have occurred simultaneously. Third, a *transmission network N* = {Ψ_1_, …, Ψ_|*N*|_} is a partition of transmission edges into disjoint transmission events. More formally, we have the following definitions.

#### Definition 3

*Given a timed phylogeny T and host labeling ℓ, an edge* (*u, v*) *of T is a transmission edge if ℓ*(*u*) ≠ *ℓ*(*v*).

#### Definition 4

*Given a timed phylogeny T and host labeling ℓ, a transmission event* Ψ *is a subset of edges of T such that (i) each edge* (*u*, *v*) ∈ Ψ *is a transmission edge, (ii) each edge* (*u*, *v*) ∈ Ψ *has the same source host ℓ*(*u*) = *s and target host ℓ*(*v*) = *t and (iii) for all pairs* (*u*, *v*), (*u*′, *v*′) ∈ Ψ *it holds that* [*τ*(*u*), *τ*(*v*)] ∩ [*τ* (*u*′); *τ* (*v*′)] ≠ ∅.

#### Definition 5

*Given a timed phylogeny T and host labeling ℓ, a transmission network N is a partition of the transmission edges of* (*T*, *ℓ*) *into disjoint transmission events*.

As suggested by the name, a transmission network *N* = {Ψ_1_, …, Ψ_|*N*|_} can be equivalently viewed as a graph. More specifically, *N* is directed, edge-labeled multi-graph, where the vertex set *V*(*N*) equals the set of hosts Σ, the edge multi-set *E*(*N*) is composed of transmission edges of *T* incurred by the host label *ℓ* associated with *N*, and the edge labeling *ψ* : *E*(*N*) → {1, …, |*N*|} assigns each transmission edge (*u*, *v*) ∈ Ψ_*i*_ to transmission event *ψ*(*ℓ*(*u*), *ℓ*(*v*))) = *i*. We say that a transmission network *N* is *consistent* with timed phylogeny *T* and host labeling *ℓ* if the set of transmission edges *N* equals the set of transmission edges in (*T*, *ℓ*).

We evaluate a transmission network *N* by two different quantities. First, the transmission number *μ*(*N*) equals the number of transmitted strain, *i.e. μ*(*N*) = Σ_Ψ∈*N*_ |Ψ|. Second, the co-transmission number *γ*(*N*) equals the number of transmission events, *i.e. γ*(*N*) = |*N*|. By definition, we have that the transmission number is greater or equal to the co-transmission number, *i.e. γ*(*N*) ≥ *μ*(*N*) for all transmission networks *N*.

Note that all transmission networks that are consistent with (*T*, *ℓ*) have the same transmission number, but may have varying co-transmission numbers. Under the principle of parsimony, we may assume that transmissions and co-transmission are rare, leading to the following optimization problem.

#### Problem 1

(*ℓ*-Transmission Network Inference (*ℓ*-TNI)) *Given a timed phylogeny T with timestamps τ and host labeling ℓ, find a transmission network N consistent with* (*T*, *ℓ*) *with minimum co-transmission number γ*(*N*).

We consider the two criteria in lexicographical order, where the first criterion seeks to minimize the number of transmitted strains, whereas the second criterion seeks to minimize the number of transmission events. Thus, we assume that the transmission of additional strains is less likely than co-transmission events by an order of magnitude. We leave exploring the trade-off between the two criteria as future work. We note that the transmission number criterion was introduced previously by Slatkin and Maddision [19], while a time-invariant version of the co-transmission number has been applied to the analyses migration in metastatic cancers [3,20]. Supplementary Table S2 provides nomenclature for topological features of transmission networks.

In practice, we do not observe a timed phylogeny *T* and host labeling *ℓ*. Rather, we obtain the genomic sequences of the strains present in individual hosts Σ. The set of extracted strains from each host forms the leaf set *L*(*T*) of an unknown timed phylogeny *T*. The function 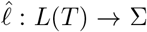 records the presence of strains in each host. As each host *s* ∈ Σ is removed from the population at time *τ*_r_(*s*), we have identical time-stamps *τ*_r_(*s*) for all strains *u* present in host *s* (*i.e.* 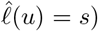. In addition, based on epidemiological data, we have an entrance time *τ*_e_(*s*) for each host *s*.

Fig. 1 shows an overview of the entities defined so far. Fig. 1a shows a timed phylogeny *T* with a leaf labeling 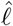 and three hosts with different entry and removal times. Figures 1b and 1c show two host labelings *ℓ*_*b*_ and *ℓ*_*c*_ respectively, both of which are consistent with the leaf labeling 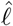. Both host labelings *ℓ*_*b*_ and *ℓ*_*c*_ have the same transmission number *μ* = 2. Further, two transmission networks *N*_*b*_ and *N*_*c*_ are shown that are consistent with the host labelings *ℓ*_*b*_ and *ℓ*_*c*_ respectively. In this case, the transmission network *N*_*c*_ has a smaller co-transmission number *γ* = 2 and is therefore more parsimonious compared to *N*_*b*_ which has a co-transmission number of *γ* = 4.

The key challenge in phylodynamics is to infer a timed phylogeny *T* and host labeling *ℓ* given leaf set *L*(*T*), host-leaf labeling 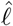, entrance times *τ*_e_ and removal times *τ*_r_. Various tools have been developed for the simpler task of inferring *T* given *L*(*T*) and *τ*_r_ [21–25]. Here, we focus on inferring a parsimonious transmission network *N* and host labeling *ℓ* given timed phylogeny *T*, host-leaf labeling 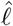, entrance times *τ*_e_ and removal times *τ*_r_.

#### Problem 2

(Transmission Network Inference (TNI)) *Given a timed phylogeny T with time-stamps τ, hostleaf labeling* 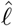, *entrance times τ_e_ and removal times τ*_r_, *find a transmission network N and corresponding host labeling ℓ with minimum transmission number μ*(*N*) = *μ*\* *and subsequently smallest co-transmission number γ*(*N*) = *γ*\* *such that τ*(*u*) ∈ [*τ*_e_(*s*), *τ*_r_(*s*)] for all hosts s and vertices *u where ℓ*(*u*) = *s*.

It is possible to define two counting versions of the above problem. The first counting problem seeks the number of transmission networks *N* with minimum transmission number *μ*(*N*) and subsequently smallest co-transmission number *γ*(*N*). The second counting problem seeks the number of host labelings *ℓ* that incur a transmission network *N* with minimum transmission number *μ*(*N*) and subsequently smallest co-transmission number *γ*(*N*). In this study, we restrict ourselves to the second version of the counting problem. Let 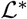 be the set of host labelings that are solutions to Problem 2. The counting problem, denoted as #TNI, is to find the cardinality of the set 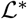 denoted by 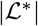. The corresponding sampling problem seeks to uniformly at random sample host labelings 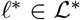.

### Complexity

The inclusion of the co-transmission number in the objective function renders the optimization and sampling versions of the TNI problem hard.

#### Complexity of the Optimization Problem

We have the following theorem.

##### Theorem 1

TNI is NP-hard.

We prove this theorem by reduction from 3-SATISFIABILITY (3-SAT), which is NP-complete [26]. In 3SAT, we are given a Boolean formula 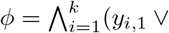 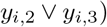with *n* variables {*x*_1_, …, *x*_*n*_} and *k* clauses in 3-conjuctive normal form (3-CNF) form. The task is to decide whether there exists a truth assignment *θ* : [*n*] → {0, 1} that satisfies all the clauses of *ϕ*. Without loss of generality, we may assume that each clause of *ϕ* consists of three distinct variables.

To relate literals to variables, we use the function *ν* : [*k*] × {1, 2, 3} → [*n*] such that *ν*(*i*, *j*) is the variable corresponding to literal *y*_*i*,*j*_. We define *σ*(*i, j*) to be 1 if *y*_*i,j*_ is a positive literal (i.e. *y*_*i,j*_ = *x*_*v*(*i,j*)_), otherwise *σ*(*i*, *j*) = 0 if *y*_*i*,*j*_ is a negative literal (*i.e.* 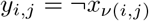). A truth assignment *θ* satisfies *ϕ* if for each clause *i* ∈ [*k*] there exists a *j* ∈ {1, 2, 3} such that *σ*(*i*, *j*) = *θ*(*ν*(*i*, *j*)).

Given *ϕ*, we construct a timed phylogeny *T*(*ϕ*) with leaf labeling 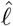 and time-stamps *τ*, *τ*_e_, *τ*_r_, as depicted in Fig. 2 and detailed below. We set 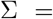 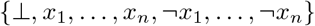. Let *ε* > 0 be a small positive constant. As for entry and removal time-stamps, we set *τ*_e_(⊥) = *τ*_r_(⊥) = 0, and 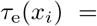 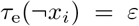 and 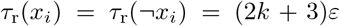 for each variable *i* ∈ [*n*]. Timed phylogeny *T*(*ϕ*) is composed of *k* clause gadgets and *n* variable gadgets, each corresponding to a subtree that is directly attached to the root *r*(*T*(*ϕ*)). The root vertex has time-stamp *τ* (*r*(*T*(*ϕ*)) = 0. The leaves of *T* have identical time-stamps (2*k* + 3)*ε*. For each variable *i* ∈ [*n*], we have a subtree *T*[var_*i*_] whose root has time-stamp *τ*(*r*(*T*[var_*i*_])) = *ε*. The two children of *r*(*T*[var_*i*_]) have identical time-stamps 2*ε*, with one child leading to two leaves labeled by positive literal *x*_*i*_ and the other child leading to two leaves labeled by negative literals 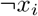. Similarly, for each clause *i* ∈ [*k*], we have a subtree *T*[clause_*i*_]. The root of this subtree has time-stamp (2*i*+1)*ε* and three children corresponding to the three literals of the clause. The three children have identical time-stamps (2*i* + 2)*ε*, each leading to two leaves labeled by the corresponding literal. Clearly, *T*(*ϕ*) can be obtained in polynomial time from *ϕ*. We refer to the supplement for the hardness proof (Supplementary Section 1.2). The supplement also shows how the reduction can be adapted to bifurcating timed phylogenies.

**Figure 2.**
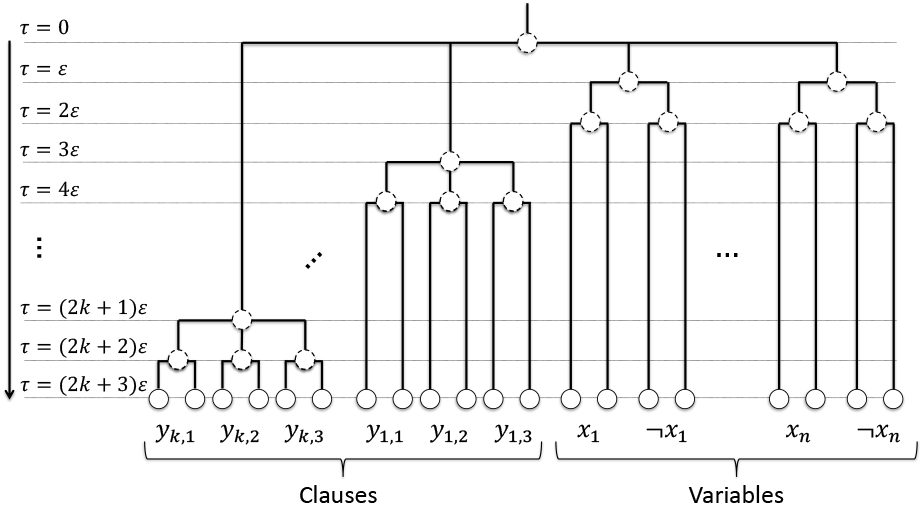
Reduction from 3-SAT to TNI. Let *ϕ* be a 3-SAT formula with *k* clauses and *n* variables. We construct a timed phylogeny *T*(*ϕ*) with vertex time-stamps indicated on the left. The host set is 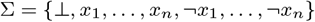. We set *τ*_e_(⊥) = *τ*_r_(⊥) = 0. For each variable *x*_*i*_ where *i* ∈ [*n*], we set 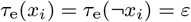 and 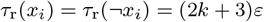. We have that *ϕ* is satisfiable if and only if there exists a minimum transmission host labeling *ℓ* of *T*(*ϕ*) with co-transmission number *γ* = 2*k* + 2*n* (Supplementary Lemma 5). Supplementary Fig. S2 shows an example.

#### Complexity of Sampling

It would be desirable to sample solutions from 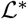, the set of host labelings *ℓ* with minimum transmission number and subsequently smallest co-transmission number, almost uniformly at random. Such a desirable algorithm is known as a *fully polynomial almost uniform sampler* (FPAUS). In general, an FPAUS for a sampling problem is a randomized algorithm that takes as input an instance *x* of the problem and a sampling tolerance *δ* > 0, and outputs a solution in time polynomial in |*x*| and log *δ*^−1^ such that the difference of the probability distribution of solutions output by the algorithm and the uniform distribution on all solutions is at most *δ* [27].

Recall the complexity class RP (randomized polynomial), which is composed of decision problems that admit randomized polynomial time algorithms that return no if the correct answer is no and otherwise return yes with probability at least 1/2. Using our reduction from 3-SAT to TNI, the existence of an FPAUS to sample the solutions of TNI would imply an FPAUS for 3-SAT. This in turn would imply that RP=NP as 3-SAT is NP-complete.

##### Theorem 2

*There exists no FPAUS to sample solutions of TNI unless RP*=*NP*.

### Simulations

To show the efficiency of our method in sampling parsimonious transmission networks, we simulate outbreaks following the procedure described in [10]. We were unable to compare to existing methods, as our simulations consider timed phylogenies which can not be used as input for joint inference methods like SCOTTI [16] and have multiple samples per host which are not supported in timed phylogeny based methods like TransPhylo [12]. However, to put the performance of our method in context we use the naive sampling algorithm as a baseline method.

We employ a two stage approach where we are given a number *m* of hosts, a transmission bottleneck size *κ* and additional epidemiological model parameters (Supplementary Section 1.5). First, we simulate a transmission process between *m* hosts using the SIR (Susceptible-Infectious-Recovered) epidemic model [28]. Under the SIR model, the outbreak begins with a single infected host and the remaining *m* – 1 individuals are infected from a unique host, each with at most *κ* co-transmitted strains. As such, the resulting transmission network *N* is a multi-tree. In the second phase, we simulate the evolution of the pathogens within each infected host using a simple coalescence model [*29*] with constant population size. Stitching together the resulting phylogenies according to *N* results in a single timed phylogeny *T*. We vary *m* ∈ {5, 10, 15, 20, 30} and *κ* ∈ {1, 2, 3}, with 5 instances for each combination, amounting to a total of 75 simulated instances. For each instance, we generate *K* = 11, 000 samples using SharpTNI and the naive sampling algorithm.

To assess the counting and sampling accuracy of our method, we restrict our attention to a subset of simulated instances (where *m* ∈ {5, 10, 15, 20} and *κ* ∈ {1, 2}) that can be exhaustively enumerated using dynamic programming (Section Methods). We find that the approximate number 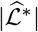 of solutions inferred by SharpTNI is nearly identical to the actual number 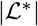 of solutions, with 69/75 instances having the correct number (Fig. 3a). Next, we compute for each solution *ℓ* in the solution set 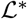, the fraction of samples generated by SharpTNI that are identical to *ℓ*. Under uniformity, this relative frequency should be close to the expected sampling frequency of 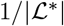. Indeed, Fig. 3b shows that the ratio between, respectively, the minimum and maximum relative frequency, and the expected sampling frequency is close to 1.

**Figure 3.**
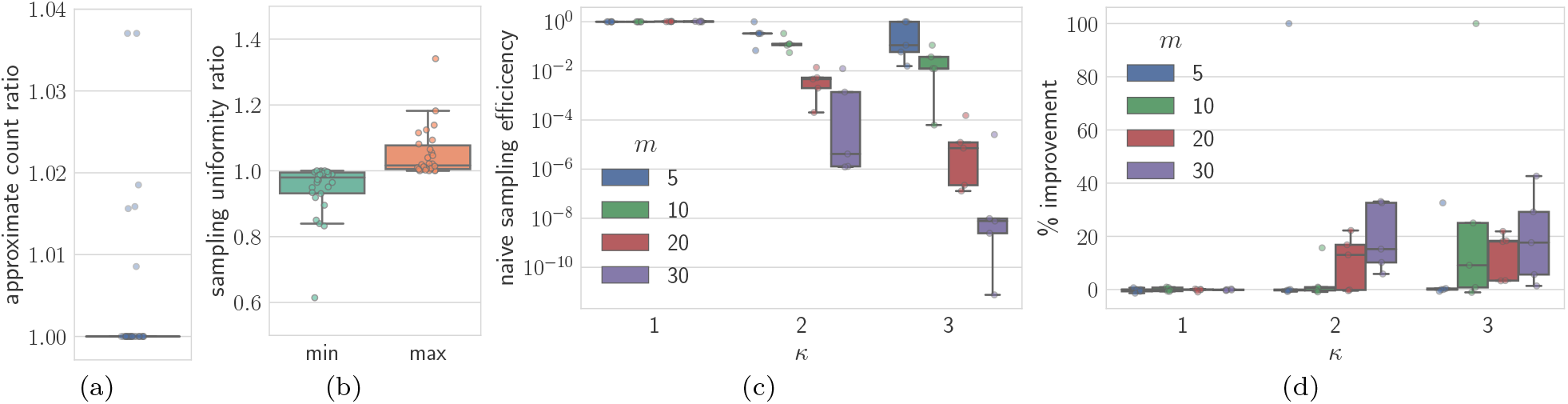
Simulations show that SharpTNI accurately counts and samples parsimonious transmission networks. Simulations were performed using a standard compartmental epidemiological model, with bottleneck size *κ* ∈ {1, 2, 3} and number *m* ∈ {5, 10, 20, 30} of hosts. SharpTNI generated *K* = 11, 000 transmission networks for each instance. (a) Ratio between the approximated number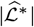 and actual number 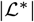 of solutions. (b) Minimum and maximum relative deviation from uniform sampling frequency 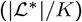. (c) Ratio between the approximate number 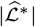 of solutions to TNI and the number 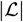 of solutions to the relaxed problem. This ratio corresponds to the success probability of the naive sampling algorithm. (d) Percentage improvement in recall of ground truth transmission edges by SharpTNI compared to the naive sampling algorithm. Supplementary Fig. S3 shows the running times.

The ratio between the number 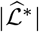 of solutions to TNI and the number of 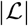 to the relaxed problem decreases exponentially with increasing number *m* of samples and the transmission bottleneck size *κ*, rendering the naive sampling algorithm impractical (Fig. 3c). Thus, we cannot expect the solutions obtained from the naive sampling algorithm to have the smallest co-transmission number *γ*. This in turn should lead to larger deviations from ground truth compared to SharpTNI. Indeed, defining *recall* as the fraction of labeled transmission edges in the ground truth host labeling *ℓ*^*^ that are correctly inferred, we observe a large relative improvement in recall by SharpTNI compared to the naive sampling algorithm (Fig. 3d). We are not showing precision, as this was identical to recall due to the ground truth transmission networks having minimum transmission number. Supplementary Fig. S3 shows the total wall time spent on a Intel Xeon 2.2 GHz processor, generating *K* = 11, 000 samples for an instance with *m* = 30 and *κ* = 3 in under 10 hours with a single thread. Since the underlying SAT sampling problem is embarrassingly parallel, SharpTNI is able to leverage UniGen’s multi-threading capabilities to cut down this running time by a factor that is equal to the number of threads.

In summary, our simulations show that SharpTNI accurately and quickly counts and samples parsimonious transmission networks, outperforming the naive sampling algorithm.

### Ebola 2014 Outbreak

To demonstrate the applicability of SharpTNI to real data, we infer parsimonious transmission networks among chiefdoms of Sierra Leone and Guinea during the 2014 Ebola outbreak [18]. The available data consist of 81 Ebola virus genomic sequences from 78 patients from Sierra Leone and 3 patients from Guinea, with metadata that include sampling date and the chiefdom where the sample was collected. There are a total of 14 Sierra Leonan chiefdoms in the data (with one chiefdom designated as unknown). Along with Guinea that makes *m* = 15 possible host labels for each node in the timed phylogeny of the 81 genomic sequences.

#### Comparison to SCOTTI

We first run SCOTTI [16], which is a Bayesian approach to co-estimate a timed phylogeny and transmission network using a Monte-Carlo Markov Chain (MCMC). We run SCOTTI for 5 × 10^6^ MCMC iterations with a burn-in percentage of 10%. We draw 100 samples of host-labeled timed phylogenies from the resulting posterior distribution. To compare the host labelings inferred by SCOTTI to those inferred by SharpTNI, we set the entry time *τ*_e_ and removal time *τ*_r_ for each host equal to the time-stamps of the first and the last node labeled by the host in that SCOTTI tree. Fig. 4a shows that the transmission numbers *μ* of the host labelings inferred by SCOTTI and SharpTNI are comparable, but that the minimum co-transmission numbers incurred by the host labelings inferred by SharpTNI are significantly smaller than those obtained using SCOTTI. This shows that SharpTNI infers a more parsimonious transmission network compared to SCOTTI.

**Figure 4.**
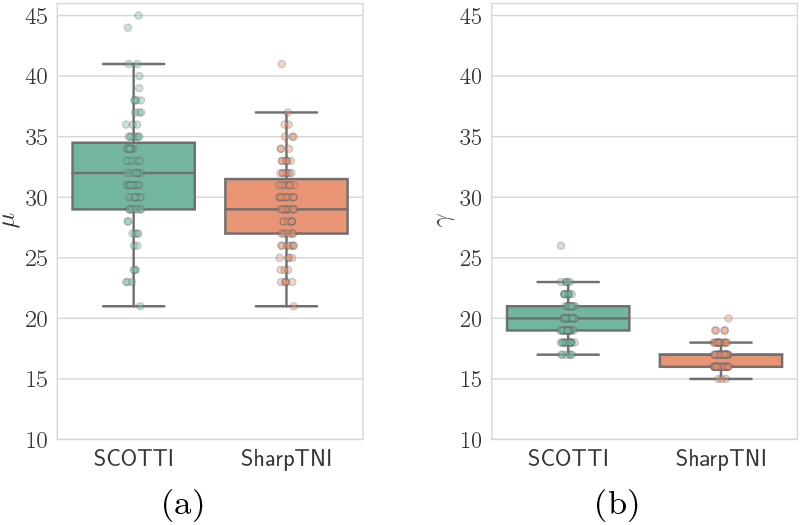
Transmission networks inferred by SharpTNI for the 2014 Ebola outbreak are more parsimonious compared to the transmission networks inferred by SCOTTI under a weak transmission bottleneck. (a) The transmission number *μ* of 100 sample trees drawn from the posterior inferred by SCOTTI are compared to the minimum transmission number inferred by SharpTNI. (b) The smallest co-transmission number *γ* of the host labeling inferred by SharpTNI is significantly smaller than the host labeling inferred by SCOTTI.

To further illustrate this point, we pick an instance where both methods inferred host labelings with the same transmission number *μ* = 24 but a co-transmission number of *γ* = 20 for SCOTTI and *γ* = 19 for SharpTNI. The transmission networks are nearly identical, except for the infection between Luawa and Jawie (Fig. 5). Notice that in both the networks, Luawa is infected by both Kissi Teng and Jawie. However, SCOTTI infers a re-infection from Luawa to Jawie whereas SharpTNI infers a transmission network with no re-infection event while keeping the transmission number the same. This leads to a simpler and more parsimonious transmission network.

**Figure 5.**
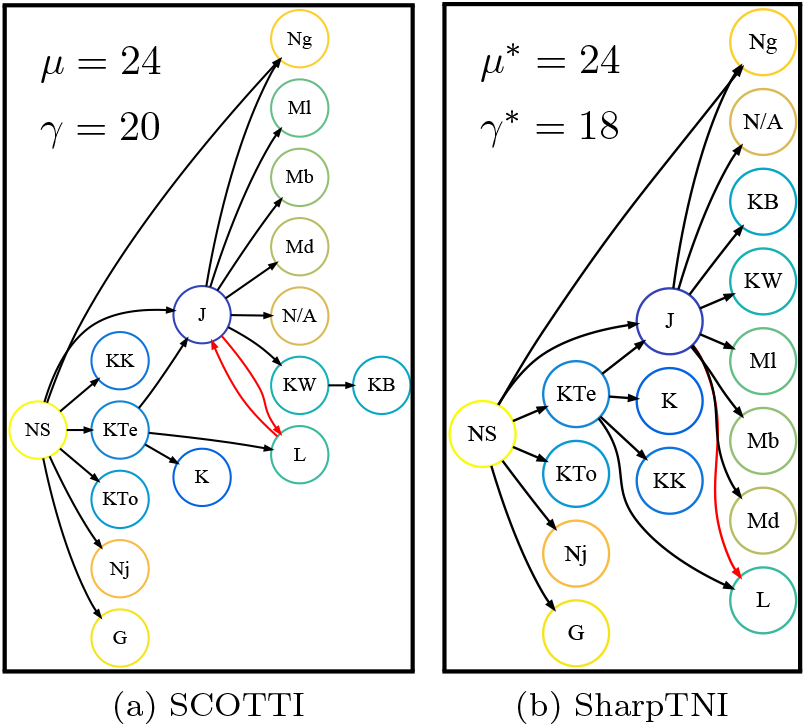
The influence of minimization of co-transmission number by SharpTNI on the inferred transmission network. (a) Transmission network inferred by SCOTTI with *μ* = 24 and *γ* = 20. (b) Transmission network inferred by SharpTNI with *μ*\* = 24 and *γ* = 18. The transmission network inferred by SCOTTI has a re-infection from Luawa to Jawie (highlighted in red) whereas in the transmission network inferred by SharpTNI there is no re-infection event. Supplementary Table S4 maps chiefdom abbreviations to full names. Supplementary Fig. S4 shows the corresponding host labelings on the timed phylogeny.

#### Re-analysis using BEAST and SharpTNI

We now re-analyze the same data using BEAST [22] to infer a timed phylogeny followed by SharpTNI to infer a transmission history. Similarly to [18], we run BEAST (version 2) for 10^6^ MCMC iterations with a burn-in percentage of 10%. Supplementary Fig. S5 shows the resulting *Maximum Clade Credibility* (MCC) consensus tree, which resembles the tree reported in [18]. We assume that a transmission from a chiefdom is possible from three weeks prior and three weeks following the first and the last sample collected from the chiefdom respectively, which is in line with reported Ebola incubation periods [30]. In addition, we allow one unsampled host in our inference with an entry and removal time that covers the entire outbreak period. Since more than 70% of the patients diagnosed in Sierra Leone were sampled, the unsampled host is most likely from Guinea. Out of a total of 324 host labelings with minimum transmission number *μ*\* = 26, SharpTNI identifies 9 transmission networks with minimum co-transmission number *γ*\* = 19 (Supplementary Fig. S6). SharpTNI identifies 9 transmission networks with minimum co-transmission number *γ*^*^ = 19 (Supplementary Fig. S6).

Gire *et al.* [18] hypothesize that the Sierra Leone outbreak stemmed from the introduction of two genetically distinct viruses from Guinea around the same time. This is because the first 12 Ebola virus disease (EVD) patients in Sierra Leone were all believed to have attended a funeral of an EVD case from Guinea and the samples from these patients fell into two distinct clusters according to their analysis. SharpTNI corroborates this hypothesis, *i*.*e*. all 9 parsimonious transmission networks (with *γ*^*^ = 19) contain a co-transmission of two strains from an unsampled host (most likely from Guinea as discussed above) to Kissi Tengi, a chiefdom located on the border of Sierra Leone and Guinea. By contrast, the majority (216/324) of host labelings that have minimum transmission number but not the smallest co-transmission number do *not* identify this co-transmission (Supplementary Fig. S7). This example highlights the utility of SharpTNI’s ability to analyze outbreaks under a weak bottleneck.

### Discussion and Conclusions

This paper introduces the Transmission Network Inference (TNI) problem for estimating a parsimonious transmission network under a weak transmission bottleneck given a timed phylogeny. Weak transmission bottlenecks arise in phylogeographic analyses of disease outbreaks as well as phylodynamics analyses of pathogens with high mutation rates, long incubation times or chronic infections. After establishing hardness of the optimization and sampling versions of the TNI problem, we present SharpTNI, a novel method for counting and sampling the solution space. The hardness of the counting problem #TNI remains open, whereas the given reduction may be used to show #P-completeness when the co-transmission number is fixed. Our method leverages recent progress in approximate counting and sampling of SATISFIABILITY [31–34]. We envision that other previously considered counting [35–39] and sampling [31,32,40] problems in computational biology can benefit similarly.

In the future, we plan to extend the current framework to co-estimation of the timed phylogeny and the transmission network by formulating a maximum likelihood version of TNI. In such a likelihood-based model, we will consider the time of transmission relative to known characteristics of the pathogen (such as incubation time). Moreover, we may assign higher likelihood to reciprocal transmissions between the same pair of hosts. In addition, we will support additional constraints such as contact maps, bottleneck sizes and other epidemiological constraints. Finally, we wish to study the problem of deriving one or more consensus transmission networks from the solution space, akin to our recent work in cancer genomics [41].

### Methods

We number the vertices of a timed phylogeny *T* from 1 to *n*, *i.e*. *V* (*T*) = {*v*_*1*_, …, *v*_*n*_}. Similarly, we number the hosts from 1 to *m*, *i.e*. Σ = {1, …, *m*}.

#### Polynomial Time Algorithm for *ℓ*-TNI

In the *ℓ*-TNI problem, we seek a transmission network *N* consistent with a given (*T, ℓ*) with minimum co-transmission number *γ*(*N*). Let *V*_*s,t*_ be a list of edges (*u, v*) of *T* where *ℓ*(*u*) = *s* and *ℓ*(*v*) = *t* sorted in ascending order by time-stamp *τ* (*v*) of the target vertex *v* (ties may be broken arbitrarily). In the following, we show that the *ℓ*-TNI problem can be reduced to 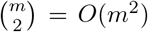 vertex partitioning problems of an interval graph, each of which can be solved by a simple greedy algorithm in time linear in |*V*_*s,t*_| [42].

For each pair (*s, t*) of distinct hosts (where *s* < *t*), we construct the interval graph *G*_*s,t*_ with vertex set *V*_*s,t*_ and an edge between (*u*, *v*) and (*u*′ *v*′) if the corresponding time intervals [*τ*(*u*); *τ*(*v*)] and [*τ*(*u*′); *τ*(*v*′)] overlap. By construction, a clique in *G*_*s,t*_ forms a set of transmission edges that can be part of the same transmission event. Thus, the minimum co-transmission number for the host pair (*s, t*) is then given by the smallest number of cliques that cover all the nodes in the interval graph. Applying the algorithm described in Ref. [42], we compute such a minimum cardinality clique partition in *O*(|*V*_*s,t*_|) time by greedily removing the maximal clique that contains the first available edge until the graph is empty (Fig. 6). Constructing the ordered sequences *V*_*s,t*_ requires *O*(*n* log *n*) time, which dominates the overall running time.

**Figure 6.**
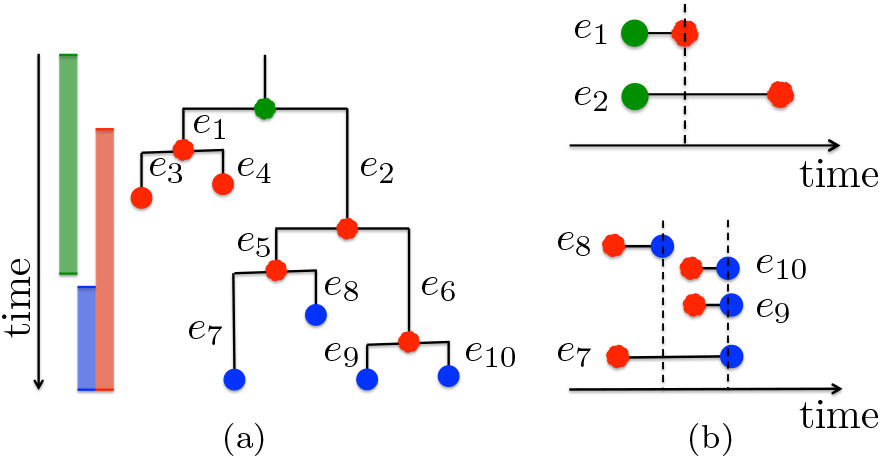
The *ℓ*-TNI problem is solvable in polynomial time. (a) An example timed phylogeny *T* with vertices colored according to a given host labeling *ℓ*. There are two sets of transmission edges, composed of two edges {*e*_1_, *e*_2_} from green to red, and four edges {*e*_7_, *e*_8_, *e*_9_, *e*_10_} from red to blue. (b) To identify a transmission network N with minimum co-transmission number *γ*(*N*) = |*N*|, we solve two clique partitioning problems on interval graphs. Given a list of transmission events between the same pair of hosts sorted by end time, we greedily identify transmission events as maximal cliques. This yields *N* = {{*e*_1_, *e*_2_}, {*e*_7_, *e*_8_}, {*e*_9_, *e*_10_}}.

#### Relaxation of TNI

To obtain a randomized algorithm for TNI, we consider a relaxation where we are interested in all host labelings *ℓ* that admit transmission networks *N* with minimum transmission number *μ*(*N*) and any co-transmission number *γ*(*N*). While the TNI problem, where we additionally require *γ*(*N*) = *γ*\*, is NP-hard, the relaxed problem can be solved in polynomial time using dynamic programming. In the following, we describe how to solve the optimization, enumeration, counting and sampling versions of this relaxed problem.

##### Optimization

Let *f*[*v, s*] be the minimum transmission number of the subtree *T*_*v*_ rooted at vertex *v* that can be attained when labeling vertex *v* by host *s, i.e. ℓ*(*v*) = *s*. The following recurrence defines *f*[*v, s*].

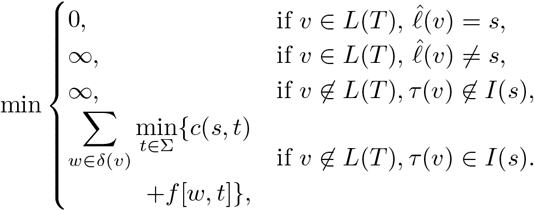

where *I*(*s*) = [*τ*_e_(*s*), *τ*_r_(*s*)], and *c*(*s, t*) = 1 if *s* = *t* and *c*(*s, t*) = 0 otherwise. The above recurrence is an adaptation of the recurrence used in the Sankoff algorithm for the small phylogeny maximum parsimony problem [43, 44]. We compute *f* bottom up from the leaves *L*(*T*) to the root vertex *r*(*T*) of *T* in *O*(*nm*) time (Supplementary Algorithm S1). The minimum transmission number *μ*\* is given by

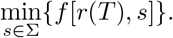

Fig. 7a shows an example of the recurrence of *f* [*v, s*] on a timed phylogeny.

**Figure 7.**
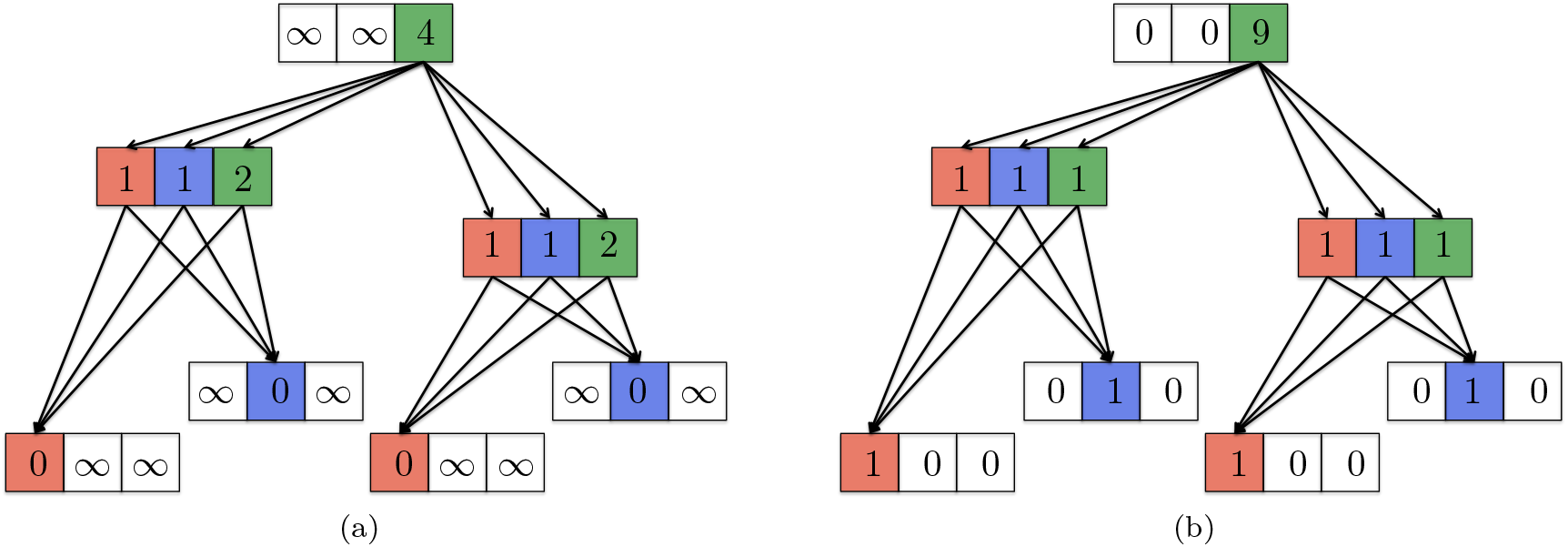
Dynamic programming for a relaxation of TNI. 4.2 example with *f, g* and *h* for the timed phylogeny shown in Fig. 1. Colored boxes (*u, s*) correspond to vertex-host pairs where *g*[*u, s*] = 1. Outgoing edges from a vertex-host pair (*u, s*) with identical target vertex *v* comprise the set Γ((*u, v*)*, s*). (a) Shows the value of *f* [*u, s*], *i.e*. the minimum transmission number of the subtree *T*_*u*_ when labeling vertex *u* with host *s*. (b) Shows the value of *h*[*u, s*], *i.e*. the number of minimum transmission labelings in the subtree *T*_*u*_ when labeling vertex *u* with host *s*.

##### Enumeration

We now identify vertex-host pairs (*v, s*) that are part of minimum transmission host labelings, indicated by *g*[*v, s*] = 1. We define *g*[*v, s*] as

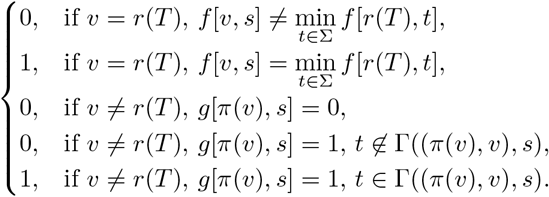

where Γ((*u, v*), *s*) is the set of host labels of vertex *v* that are part of minimum transmission host labelings *ℓ* where the parent vertex *u* is labeled by host *s*, *i.e*. Γ((*u, v*), *s*) = {*t* ∈ Σ | *c*(*s, t*) + *f* [*v, t*] = min_*t*′∈Σ_{*c*(*s, t*′)+*f*[*v, t*′]}}. We note that *g* can be computed in a top down fashion in *O*(*nm*) time (Supplementary Algorithm S2), whereas Γ can be computed in *O*(*m*) time. Using *g* and Γ, we enumerate all minimum transmission host labelings of *T* (Supplementary Algorithm S3 and S4).

##### Counting

Next, we consider the counting version of this problem. This number can also be solved using dynamic programming [45]. Let *h*[*v, s*] denote the number of minimum transmission labelings in the subtree *T*_*v*_ of *T* rooted at vertex *v* when *ℓ*(*v*) = *s*. We define *h*[*v, s*] recursively as

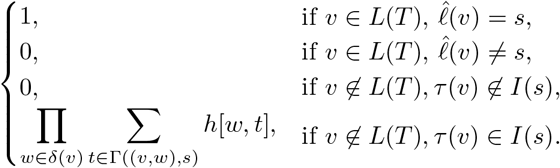

The total number of solutions is given by

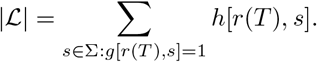

Directly translating the above recurrence into a recursive function results in a *O*(*nm*) time algorithm. Fig. 7b shows an example of the recurrence of *h*[*v, s*] on a timed phylogeny.

##### Sampling

Using the count matrix *h*[*u, s*], we introduce a subroutine that takes a vertex *v* and host *s* as input, and uniformly samples a host labeling *ℓ_u_* of subtree *T*_*u*_ rooted at *u* subject to the restriction that *ℓ*_*u*_(*u*) = *s* (Supplementary Algorithm S5). Supplementary Section 1.3 gives a correctness proof of our algorithm.

Let Σ* = {*s*_1_, …, *s*_*k*_} be the set of hosts of the root vertex *r*(*T*) that are part of minimum transmission labelings, *i.e*. Σ* = {*s* ∈ Σ | *g*[*r*(*T*), *s*] = 1}. The fraction *p*_*s*_ of minimum transmission host labelings *ℓ* where *ℓ*(*r*(*T*)) = *s* equals *h*[*r*(*T*), *s*]/ Σ_*s*′Σ*_ *h*[*r*(*T*), *s*′]. Thus, to sample *all* minimum transmission host labelings uniformly at random, we draw a *s* ∈ Σ* according to the categorical probability distribution defined by (*p*_1_, …, *p*_*k*_). Supplementary Algorithm S6 is then used on *T* with *ℓ*(*r*(*T*)) = *s* to sample minimum transmission host labeling *ℓ* of *T* uniformly at random. This takes *O*(*nm*) time per sample.

##### Naive sampling algorithm

To identify host labelings with minimum transmission number and subsequently smallest co-transmission number, we may repeatedly generate a uniformly random sample using the above algorithm and retain only those host labelings that have smallest co-transmission number. The success probability of this naive sampling algorithm is 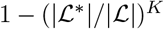 where *K* is the number of repetitions.

#### Solving TNI via SAT

We focus our attention on a decision version of the general TNI problem: is there a host labeling *ℓ* that admits a transmission network *N* with transmission number *μ*(*N*) = *μ*\* and co-transmission number *γ*(*N*) = *α*, where *α* ∈ ℕ? Since *γ*\* ∈ {|Σ| − 1, …, |*E*|}, we may solve the optimization problem of finding *N* with minimum *γ*(*N*) = *γ*\* by initially setting *α* = |Σ| − 1 = *m* − 1 and incrementing *α* until the decision problem has a yes-answer or *α* = |*E*(*T*)| = *n* − 1.

In the following, we will show how to reduce a TNI instance 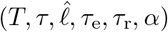 to a Boolean formula *ϕ*. To facilitate almost uniform sampling and approximate counting, we require that there is a bijection between the solutions to TNI instance 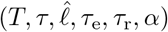 and the corresponding SAT instance *ϕ*. As such, we must introduce variables and constraints that encode (i) a host labeling *ℓ*, (ii) *ℓ* has minimum transmission number *μ*\*, (iii) *ℓ* admits a transmission network *N* with co-transmission number *γ*(*N*) = *α* and (iv) uniqueness of *N* given *ℓ*.

For clarity, we will not present constraints in clause normal form (CNF). Rather, we refer the reader to Supplementary Section 1.4 for a CNF representation of *ϕ* with *O*(*n*^2^ + *nm* + *nα*) variables and *O*(*nm*^2^*α*^2^ + *n*^2^*m*^2^ + *n*^2^*α*^2^) clauses.

##### Host labeling

Variables **x** ∈ {0, 1}^*n*×*m*^ encode a host labeling. That is *x*_*i,s*_ = 1 if vertex *v*_*i*_ is labeled by host *ℓ*(*v*_*i*_) = *s*, and *x*_*i,s*_ = 0 otherwise. To encode a host labeling, we introduce the following constraints for all vertices *v*_*i*_ ∈ *V* (*T*).

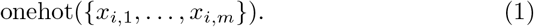

The function onehot(*X*) encodes that exactly one binary variable *x* ∈ *X* is true, which can be accomplished by the following constraint.

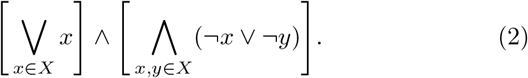

##### Minimum transmission number μ*

Next, we need to ensure that **x** encodes a host labeling with minimum transmission number *μ*\*. To this end, we use the functions *f*, *g* and Γ defined in the previous section. First, we prevent labeling a vertex *v*_*i*_ by a host *s* if this is not part of a minimum transmission host labeling (*i.e*., *g*(*v*_*i*_, *s*) = 0). That is, for all vertex-host pairs (*v*_*i*_, *s*) ∈ *V* (*T*) × [*m*] where *g*[*v*_*i*_, *s*] = 0, we have

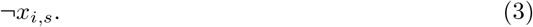

Labeling a vertex *v*_*i*_ by host *s* restricts the set of host for each child *v_j_* of *v*_*i*_ to Γ((*v*_*i*_, *v*_*j*_), *s*). Thus, for all edges (*v*_*i*_, *v*_*j*_) ∈ *E*(*T*) and hosts *s* ∈ [*m*], we have

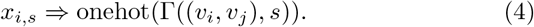

##### Transmission network

We now need to encode that **x** admits a transmission network *N* with cotransmission number *γ*(*N*) = *α*. We order the edges *E*(*T*) = {*e*_1_, …, *e*_*n*−1_} in ascending order by the time-stamp of the target vertex, breaking ties arbitrarily. We introduce a variable *c*_*ij,kl*_ for each pair (*i, j*), (*k, l*) of distinct edges where (*i, j*) < (*k, l*). We require *c*_*ij,kl*_ = 1 if and only if (*i, j*) and (*k, l*) are transmission edges between the same pair of hosts with overlapping time intervals. This is achieved by the following three sets of constraints. First, we have

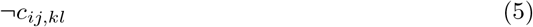

for all edge pairs (*i, j*) < (*k, l*) that do not have overlapping time intervals, *i.e*. [*τ*(*v*_*i*_), *τ*(*v*_*j*_)] ∩ [*τ*(*v*_*k*_), *τ*(*v*_*l*_)] = ∅. Second, we have that *c*_*ij,kl*_ = 0 for all edges (*i, j*) < (*k, l*) where *ℓ*(*v*_*i*_) = *ℓ*(*v*_*j*_) or *ℓ*(*v*_*k*_) = *ℓ*(*v*_*l*_). That is, for all edge pairs (*i, j*) < (*k, l*) and hosts *s* ∈ [*m*], we have

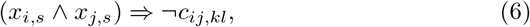

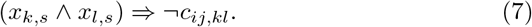

Third, *c*_*ij,kl*_ = 1 if (*i, j*) and (*k, l*) are transmission edges between the same pair of hosts with overlapping time intervals. That is, for all pairs (*i, j*) < (*k, l*) of distinct edges with overlapping time intervals, *i.e*. [*τ*(*v*_*i*_), *τ*(*v*_*j*_)] ∩ [*τ*(*v*_*k*_), *τ*(*v*_*l*_)] = ∅ and hosts *s,t* ∈ [*m*] where 1 < *s* < *t* < *m* we have

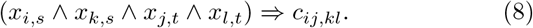

We now introduce variables **y** ∈ {0, 1}^(*n*−1)×*α*^ such that *y*_*ij,p*_ = 1 if and only if (*i, j*) is a transmission edge and assigned to transmission event *p*. We require that each transmission edge (*i, j*) is assigned to exactly one transmission event. That is, for all edges (*i, j*) and distinct hosts *s* < *t*, we have

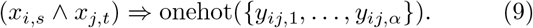

Next, if (*i, j*) is not a transmission edge then it must not be assigned to any transmission event *p*. That is, for all edges (*i, j*), hosts *s* and transmission events *p* ∈ [*α*], we have

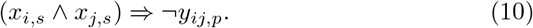

Finally, edges (*i, j*) < (*k, l*) that are not time-overlapping, transmission edges between the same pair of hosts (*i.e*. *c*_*ij,kl*_ = 0), must not be assigned to the same transmission event *p* ∈ [*α*]. That is, for all distinct edges (*i, j*) < (*k, l*) and transmission events *p* ∈ [*α*], we have

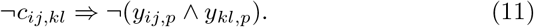

##### Uniqueness

To ensure bijectivity between the set of satisfying assignments of *ϕ* and the set of host labelings *ℓ* that admit a transmission network *N* with transmission number *μ*(*N*) = *μ*\* and co-transmission number *γ*(*N*) = *α*, we require that each host labeling *ℓ* encodes a unique transmission network *N*. To that end, we introduce constraints that will pick the exact same transmission network *N* given *ℓ* as the greedy algorithm described in Section Methods. Specifically, each transmission edge (*k, l*) must be assigned to the same transmission event *p* as the first transmission edge (*i, j*) that overlaps in time and hosts with (*k, l*) (*i.e*. *c_ij,kl_* = 1). That is, for all edges (*i, j*) < (*k, l*) and transmission events *p* ∈ [*α*], we have

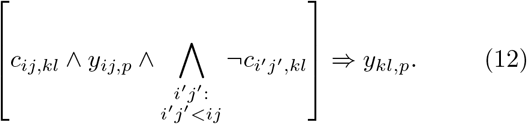

While variables **x** uniquely determine variables **c**, they do not uniquely determine variables **y** as there exist *α*! permutations of the *α* transmission events. To break this symmetry, we use the edge ordering *E*(*T*) = {*e*_1_, …, *e*_*n*−1_} to designate the smallest transmission edge of each transmission event *p* as its representative. We require that these representatives are assigned to transmission events according to the edge ordering. Specifically, we introduce variables **z** ∈ {0, 1}^*n*−1^ such that *z*_*ij*_ = 1 if and only if edge (*i, j*) is a representative transmission edge of some transmission event.

We impose the forward direction of the bi-implication by modeling the contrapositive using the following two set of constraints. First, if edges (*i, j*) < (*k, l*) are assigned to the same transmission event *p* then edge (*k, l*) cannot be a representative. That is, for all distinct edges (*i, j*) < (*k, l*) and transmission events *p* ∈ [*α*], we have

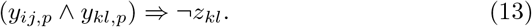

Second, if an edge (*i, j*) is not a transmission edge then it cannot be a representative. That is, for all edges (*i, j*) and hosts *s*, we have

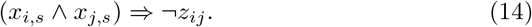

To model the reverse direction, we have for all edges (*i, j*) < (*k, l*) and transmission events *p* ∈ [*α*]

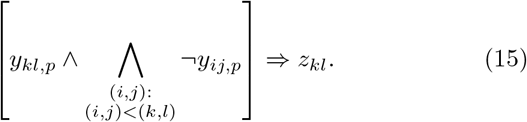

Finally, we require that representatives are ordered correctly. For all representatives (*i, j*) < (*k, l*) where (*i, j*) is assigned to transmission event *q*, it cannot be that (*k, l*) is assigned to a transmission event *p* < *q*. That is, for all edges (*i, j*) < (*k, l*) and transmission events *p* < *q*, we have

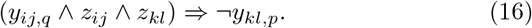

##### Approximate counting and almost uniform sampling

Now that we have a SAT formula, we look at the related problems of approximate sampling and almost uniform sampling of the solution space [46]. We use ApproxMC [33, 34] to approximate 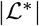 and Uni-Gen [31, 32] to sample almost uniformly from 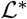. We call the resulting method SharpTNI.

## List of Abbreviations

TNI: Transmission Network Inference
SAT: Satisfiability
CNF: Conjunctive Normal Form
FPAUS: Fully Polynomial Almost Uniform Sampler
RP: Randomized Polynomial
NP: Nondeterministic Polynomial
SIR: Susceptible-Infectious-Recovered
MCC: Maximum Clade Crediblity
EVD: Ebola Virus Disease

## Declarations

## Author’s contributions

M.E-K. conceived the project. P.S. developed the code and performed the experimental evaluation. P.S. and M.E-K. wrote the manuscript. All authors read and approved the final manuscript.

## Acknowledgements

M.E-K. was supported by the National Science Foundation (CCF 18-50502). Experiments were run on Blue Waters, which is a joint effort of the University of Illinois at Urbana-Champaign and the National Center for Supercomputing Applications. The authors thank the anonymous referees for insightful comments that have improved the manuscript.

## Consent for publication

Not applicable.

## Ethics approval and consent to participate

Not applicable.

## Availability of data and materials

Ebola and simulated data used in the results section is available at https://github.com/elkebir-group/SharpTNI/tree/master/data. Results generated using this data are available at https://doi.org/10.13012/B2IDB-9734610_V1.

## Supplementary Materials

Background and Problem Statement — Fig. S1, Tables S1 and S2

Complexity Proof — Lemmas 1 to 4

Algorithms for Relaxed TNI Problem — Algorithms S1 to S6

CNF form of the SAT formulation — Eqs. 2 to 19, Table S3

Outbreak Simulation Details — Text

Simulated and Ebola Outbreak Analysis — Figs. S3 to S7, Table S4

## References

1. Dellicour, S., Baele, G., Dudas, G., Faria, N.R., Pybus, O.G., Suchard, M.A., Rambaut, A., Lemey, P.: Phylodynamic assessment of intervention strategies for the West African Ebola virus outbreak. Nature communications 9(1), 2222 (2018)

2. Romero-Severson, E., Skar, H., Bulla, I., Albert, J., Leitner, T.: Timing and order of transmission events is not directly reflected in a pathogen phylogeny. Molecular biology and evolution 31(9), 2472–2482 (2014)

3. El-Kebir, M., Satas, G., Raphael, B.J.: Inferring parsimonious migration histories for metastatic cancers. Nature Genetics 50(5), 718–726 (2018)

4. Leitner, T., Escanilla, D., Franzen, C., Uhlen, M., Albert, J.: Accurate reconstruction of a known HIV-1 transmission history by phylogenetic tree analysis. Proceedings of the National Academy of Sciences 93(20), 10864–10869 (1996)

5. Cottam, E.M., Thébaud, G., Wadsworth, J., Gloster, J., Mansley, L., Paton, D.J., King, D.P., Haydon, D.T.: Integrating genetic and epidemiological data to determine transmission pathways of foot-and-mouth disease virus. Proceedings of the Royal Society B: Biological Sciences 275(1637), 887–895 (2008)

6. Harris, S.R., Feil, E.J., Holden, M.T., Quail, M.A., Nickerson, E.K., Chantratita, N., Gardete, S., Tavares, A., Day, N., Lindsay, J.A., et al.: Evolution of mrsa during hospital transmission and intercontinental spread. Science 327(5964), 469–474 (2010)

7. Ypma, R.J., Bataille, A., Stegeman, A., Koch, G., Wallinga, J., Van Ballegooijen, W.M.: Unravelling transmission trees of infectious diseases by combining genetic and epidemiological data. Proceedings of the Royal Society B: Biological Sciences 279(1728), 444–450 (2011)

8. Snitkin, E.S., Zelazny, A.M., Thomas, P.J., Stock, F., Henderson, D.K., Palmore, T.N., Segre, J.A., Program, N.C.S., et al.: Tracking a hospital outbreak of carbapenem-resistant klebsiella pneumoniae with whole-genome sequencing. Science translational medicine 4(148), 148–116148116 (2012)

9. Ypma, R.J., van Ballegooijen, W.M., Wallinga, J.: Relating phylogenetic trees to transmission trees of infectious disease outbreaks. Genetics 195(3), 1055–1062 (2013)

10. Didelot, X., Gardy, J., Colijn, C.: Bayesian inference of infectious disease transmission from whole-genome sequence data. Molecular biology and evolution 31(7), 1869–1879 (2014)

11. Hall, M., Woolhouse, M., Rambaut, A.: Epidemic reconstruction in a phylogenetics framework: transmission trees as partitions of the node set. PLoS computational biology 11(12), 1004613 (2015)

12. Didelot, X., Fraser, C., Gardy, J., Colijn, C.: Genomic infectious disease epidemiology in partially sampled and ongoing outbreaks. Molecular biology and evolution 34(4), 997–1007 (2017)

13. Klinkenberg, D., Backer, J.A., Didelot, X., Colijn, C., Wallinga, J.: Simultaneous inference of phylogenetic and transmission trees in infectious disease outbreaks. PLoS computational biology 13(5), 1005495 (2017)

14. Skums, P., Zelikovsky, A., Singh, R., Gussler, W., Dimitrova, Z., Knyazev, S., Mandric, I., Ramachandran, S., Campo, D., Jha, D., et al.: QUENTIN: reconstruction of disease transmissions from viral quasispecies genomic data. Bioinformatics 34(1), 163–170 (2017)

15. Leonard, A.S., Weissman, D.B., Greenbaum, B., Ghedin, E., Koelle, K.: Transmission bottleneck size estimation from pathogen deep-sequencing data, with an application to human influenza A virus. Journal of virology 91(14), 00171–17 (2017)

16. De Maio, N., Wu, C.-H., Wilson, D.J.: Scotti: efficient reconstruction of transmission within outbreaks with the structured coalescent. PLoS computational biology 12(9), 1005130 (2016)

17. De Maio, N., Worby, C.J., Wilson, D.J., Stoesser, N.: Bayesian reconstruction of transmission within outbreaks using genomic variants. PLoS computational biology 14(4), 1006117 (2018)

18. Gire, S.K., Goba, A., Andersen, K.G., Sealfon, R.S., Park, D.J., Kanneh, L., Jalloh, S., Momoh, M., Fullah, M., Dudas, G., et al.: Genomic surveillance elucidates ebola virus origin and transmission during the 2014 outbreak. science 345(6202), 1369–1372 (2014)

19. Slatkin, M., Maddison, W.P.: A cladistic measure of gene flow inferred from the phylogenies of alleles. Genetics 123(3), 603–613 (1989)

20. El-Kebir, M.: Parsimonious migration history problem: Complexity and algorithms. In: 18th International Workshop on Algorithms in Bioinformatics, WABI 2018, August 20-22, 2018, Helsinki, Finland, pp. 24–12414 (2018). doi:10.4230/LIPIcs.WABI.2018.24

21. Drummond, A.J., Rambaut, A.: BEAST: Bayesian evolutionary analysis by sampling trees. BMC evolutionary biology 7(1), 214 (2007)

22. Bouckaert, R., Heled, J., Kühnert, D., Vaughan, T., Wu, C.-H., Xie, D., Suchard, M.A., Rambaut, A., Drummond, A.J.: BEAST 2: a software platform for bayesian evolutionary analysis. PLoS computational biology 10(4), 1003537 (2014)

23. Stamatakis, A.: RAxML version 8: a tool for phylogenetic analysis and post-analysis of large phylogenies. Bioinformatics 30(9), 1312–1313 (2014)

24. Price, M.N., Dehal, P.S., Arkin, A.P.: FastTree 2–approximately maximum-likelihood trees for large alignments. PloS one 5(3), 9490 (2010)

25. Bouckaert, R., Vaughan, T.G., Barido-Sottani, J., Duchêne, S., Fourment, M., Gavryushkina, A., Heled, J., Jones, G., Kühnert, D., De Maio, N., et al.: BEAST 2.5: An advanced software platform for bayesian evolutionary analysis. PLoS computational biology 15(4), 1006650 (2019)

26. Karp, R.M.: In: Miller, R.E., Thatcher, J.W., Bohlinger, J.D. (eds.) Reducibility among Combinatorial Problems, pp. 85–103. Springer, Berlin, Heidelberg (1972)

27. Jerrum, M.: Counting, Sampling and Integrating: Algorithms and Complexity. Springer, Berlin, Heidelberg (2003)

28. Allen, L.J.: An introduction to stochastic epidemic models. In: Mathematical Epidemiology, pp. 81–130. Springer, Berlin, Heidelberg (2008)

29. Kingman, J.: b the coalescent. stoch. In: Proc. Appl, vol. 13, pp. 235–248 (1982)

30. Eichner, M., Dowell, S.F., Firese, N.: Incubation period of ebola hemorrhagic virus subtype zaire. Osong public health and research perspectives 2(1), 3–7 (2011)

31. Chakraborty, S., Meel, K.S., Vardi, M.Y.: Balancing scalability and uniformity in sat witness generator. In: Proceedings of the 51st Annual Design Automation Conference, pp. 1–6 (2014). ACM

32. Chakraborty, S., Fremont, D.J., Meel, K.S., Seshia, S.A., Vardi, M.Y.: On parallel scalable uniform sat witness generation. In: International Conference on Tools and Algorithms for the Construction and Analysis of Systems, pp. 304–319 (2015). Springer

33. Chakraborty, S., Meel, K.S., Vardi, M.Y.: A Scalable Approximate Model Counter. In: Principles and Practice of Constraint Programming, pp. 200–216. Springer, Berlin, Heidelberg (2013)

34. Soos, M., Meel, K.S.: BIRD: Engineering an efficient CNF-XOR SAT solver and its applications to approximate model counting. In: Proceedings of AAAI Conference on Artificial Intelligence (AAAI)(1 2019) (2019)

35. Miklós, I., Kiss, S.Z., Tannier, E.: Counting and sampling SCJ small parsimony solutions. Theoretical Computer Science 552, 83–98 (2014)

36. Chauve, C., Courtiel, J., Ponty, Y.: Counting, generating, analyzing and sampling tree alignments. International Journal of Foundations of Computer Science 29(05), 741–767 (2018)

37. Chauve, C., Ponty, Y., Wallner, M.: Counting and sampling gene family evolutionary histories in the duplication-loss and duplication-loss-transfer models. arXiv preprint arXiv:1905.04971 (2019)

38. Ponty, Y.: Efficient sampling of RNA secondary structures from the boltzmann ensemble of low-energy. Journal of mathematical biology 56(1–2), 107–127 (2008)

39. Dyer, M.: Approximate counting by dynamic programming. In: Proceedings of the Thirty-fifth Annual ACM Symposium on Theory of Computing, pp. 693–699 (2003). ACM

40. Bellare, M., Goldreich, O., Petrank, E.: Uniform generation of NP-witnesses using an NP-oracle. Information and Computation 163(2), 510–526 (2000)

41. Aguse, N., Qi, Y., El-Kebir, M.: Summarizing the solution space in tumor phylogeny inference using multiple consensus trees. Bioinformatics (ISMB/ECCB 2019) In press (2019)

42. Finke, G., Jost, V., Queyranne, M., Sebő, A.: Batch processing with interval graph compatibilities between tasks. Discrete Applied Mathematics 156(5), 556–568 (2008)

43. Sankoff, D.: Minimal mutation trees of sequences. SIAM Journal on Applied Mathematics 28(1), 35–42 (1975)

44. Fitch, W.M.: Toward defining the course of evolution: minimum change for a specific tree topology. Systematic Biology 20(4), 406–416 (1971)

45. Giegerich, R., Meyer, C.: Algebraic dynamic programming. In: International Conference on Algebraic Methodology and Software Technology, pp. 349–364 (2002). Springer

46. Jerrum, M.R., Valiant, L.G., Vazirani, V.V.: Random generation of combinatorial structures from a uniform distribution. Theoretical Computer Science 43, 169–188 (1986)

